# Linguistic and Affective Prosody – A Unifying Perspective: A Systematic Review and ALE Meta-Analysis of the Cortical Organization of Prosody

**DOI:** 10.1101/2024.10.29.620829

**Authors:** Charalambos Themistocleous

**Affiliations:** Department of Special Needs Education, University of Oslo, Oslo, Norway

## Abstract

Prosody is a cover term referring to the melodic aspects of speech, with linguistic and affective (a.k.a. emotional) meanings. This review provides an overview of linguistic and affective prosody, testing two hypotheses on healthy individuals’ linguistic and affective prosody. The first hypothesizes that the biological nature of affective prosody triggers activations unrelated to language (biological hypothesis), and the second that the aspects of affective prosody have been grammaticalized, i.e., incorporated into the language (linguistic hypothesis). We employed a systematic ALE metanalytic approach to identify neural correlates of prosody from the literature. Specifically, we assessed papers that report brain coordinates from healthy individuals selected using systematic research from academic databases, such as PubMed (NLM), Scopus, and Web of Science. We found that affective and linguistic prosody activate bilateral frontotemporal regions, like the Superior Temporal Gyrus (STG). A key difference is that affective prosody involves subcortical structures like the amygdala, and linguistic prosody activates linguistic areas and brain areas of social cognition and engagement. The shared activations, therefore, suggest that linguistic and affective meanings are combined, involving shared underlying brain connectivity mechanisms and acoustic manifestations. We propose the “prosodic blending hypothesis” to account for the findings. The hypothesis unifies the linguistic and affective prosody distinction, suggesting that prosody incorporates affective, social, and linguistic meanings in the acoustic signal and involves selective bilateral neural contributions. Although brain regions responsible for core emotions can be activated, those meanings are produced through a coordinated and structured linguistic manner, accounting for the gradient expression of non-linguistic and categorical linguistic aspects. We argue that the process is similar to speech, lexicon, and grammar domains, as they, too, convey affective, social, and linguistic meanings without their explicit separation into linguistic and affective. Therefore, our blending hypothesis views prosody as a unified system that combines affective and linguistic functions.

## Introduction

Research on the neurocognitive foundations of prosody has essential theoretical and practical implications and has proliferated in the last forty years. This review focuses on the brain processes involved in prosodic production. Prosody is the ability to produce and perceive the melodic patterns of speech and their linguistic or non-linguistic meanings. It is a cover term involving several melodic properties, like pitch, loudness, quantity, speech rate, and voice quality features, such as aspiration and nasalization (Lehiste, 1970).

In clinical research, there has been a twofold distinction between affective (a.k.a., emotional) and linguistic functions of prosody (Gussenhoven, 2004b; Ladd, 2009). Affective prosody expressing anger, joy, affection, and sadness is typically language-independent and spans diverse societies and cultures (Cordaro et al., 2018). On the other hand, linguistic prosody is considered a component of the linguistic system; being part of grammar, it interfaces with language domains, such as phonology, morphology, syntax, semantics, and pragmatics (Erteschik-Shir, 2007; Lambrecht, 1994; Zubizarreta, 1998). Typically, linguistic prosody is distinguished into lexical prosody and post-lexical prosody. Lexical prosody concerns lexical prominence, such as the lexical accent, stress, tone, and temporal properties, like syllable quantity (Gordon, 2006). Based on the lexical prominence, languages are traditionally classified into pitch accent languages, like Norwegian and Serbo-Croatian; stress languages, like English and Greek; and tonal languages, like Chinese and Vietnamese (see also, Jun (2010)).

Post-lexical prosody spans over one or more words (Lehiste, 1970). It highlights phonological constituents (e.g., words and phrases), marks their boundaries, and assigns melodic patterns to the utterance, such as melodies in polar and wh-questions, statements, and commands. Intonation plays a central role in post-lexical prosody and has been the subject of considerable research (’t Hart et al., 1990; Bolinger, 1947; Bruce, 1977; Halliday, 1967; Haugen & Joos, 1952/1972; Liberman, 1979; Palmer, 1922; J. Pierrehumbert, 1980; Pike, 1945; Xu, 1996). Intonation typically refers to the arrangement of the melodic units of an utterance to create well-formed melodic patterns (Arvaniti & Ladd, 1995; Beckman & Pierrehumbert, 1986b; J. B. Pierrehumbert, 1980; Prieto, 2005; Xu, 1998; Yu, 2007).

Although prosody is strongly linked to specialized brain areas for prosody, the details of this relationship remain controversial. One source of controversy involves the conflicting evidence about the lateralization of prosody, as substantial evidence supports the involvement of mostly right-lateralized brain areas in prosody (Lancker & Breitenstein, 2000). Evidence from patients with prosody impairments linked to emotional expression, such as monotonous speech, can be indicative of neurological conditions like Alzheimer’s disease, Parkinson’s disease, and Primary Progressive Aphasia (J. D. Rohrer et al., 2012), provides further support from the hemispheric specialization of prosody. For example, findings from patients with RH stroke show impairments in affective prosody (Durfee et al., 2021; Sheppard et al., 2021; Wright et al., 2018). However, others reported insignificant functional lateralization for emotional prosody perception (Witteman et al., 2014).

Another critical concern is how affective and linguistic prosody interface with specific brain regions of interest (ROIs)—that is, whether affective prosody activates distinct ROIs not involved in linguistic prosody. Affective prosody is linked to the expression of emotions, a core biological function in humans and animals. This biological foundation can explain the universal prosodic characteristics of emotions across cultures and languages (Ekman et al., 2013). However, since humans express emotions through conventional linguistic and social means, ROIs associated with both emotion and language can be involved. Moreover, emotions interpret the linguistic message, providing different connotations, such as when expressing judgment or irony (Jing-Schmidt & Kapatsinski, 2012; Panksepp, 2005). To account for the complex functions of prosody and how it activates different areas, this study proposes two hypotheses: the biological hypothesis and the linguistic hypothesis. The *biological hypothesis* suggests that affective and linguistic prosody are processed by distinct brain areas, with affective prosody tied to emotional expression in social and biological contexts, and linguistic prosody linked to language processing. However, in humans, while affective prosody may serve a biological function as in other animals (e.g., signaling aggression or submission), it has also been adapted to fit social conventions, as seen in the use of prosody to convey subtle social cues (Eckert, 2019; Foulkes & Docherty, 2006; Silverstein, 2010). The *linguistic hypothesis* is an extension of this idea and proposes that affective prosody can be integrated into the linguistic system through a *grammaticalization* process where aspects of speech become more conventionalized and systematic. This process of grammaticalization, as proposed by (Gussenhoven, 2002, 2004a), implies that emotional vocalizations have acquired conventional linguistic functions, thus following the standard linguistic pathways of speech perception, such as the dual-stream model (Hickok & Poeppel, 2007).^1^

To account for the grammaticalization of prosody, Gussenhoven (2002, 2004a) proposes three codes, the *frequency code*, which predicts that larger and more aggressive animals are producing lower pitch whereas smaller animals produce higher pitch productions, and these were grammaticalized in human language so that higher pitch was grammaticalized to indicate politeness and express questions and lower pitch definiteness and completion of utterances (Gussenhoven, 2002, 2004a). The *effort code* suggests that deliberate and precise control of pitch has been grammaticalized to express emphasis (Gussenhoven, 2002, 2004a): pitch accents that highlight specific parts of the utterance are demarcating the focused parts more precisely than the non-emphatic pitch movement has grater window of flexible alignment often overshooting or undershooting (Themistocleous, 2016b). Lastly, the *production code* is associated with the control of how fundamental frequency declines during the production of the utterance. The phenomenon known as declination is a physiological process where the F0 typically starts at higher frequency and ends in a lower one because of energy exhaustion over the course of an utterance. The grammaticalized control of declination allows speakers to reverse the typical *F0* pattern and raise the *F0* at the right boundary of a phrase to signal their intention to continue the utterance (alternatively, an *F0* fall at the right boundary can function as signal that allows another speaker to take the floor of the conversation) (Sacks et al., 1974).

In this review paper, we concentrate on trying to systematize the large literature that has examined the degree of cortical areas and different measures of prosody and prosodic functions and focus on two critical theoretical issues: (1) the nature of explicit prosodic brain areas and (2) the distinction between linguistic and affective prosody. Thus, the primary questions are (1) whether the affective and linguistic prosody are associated with distinct neural correlates and (2) whether the two types of prosody are part of two different systems as suggested by the biological and linguistic hypothesis or simply two sides of the same coin. To answer this question, we will provide in the following part a rather lengthy and comprehensive overview of the nature of linguistic and affective prosody based on well-received views on prosody to equip the reader with a more in-depth understanding of the characteristics and differences in linguistic and affective prosody. Subsequently, we will explore those characteristics and differences through an ALE meta-analysis, where we evaluate the two hypotheses biological and linguistic hypotheses on the relationship between prosodic and affective prosody. After presenting a systematic ALE meta-analytic review of the evidence from correlational studies, we return to questions about the possible causal status of the effects demonstrated in the discussion. We believe that this relational literature has important implications for the causal hypotheses on the link between prosody and the brain and provides a perspective about impaired prosody, which is, in turn, a critical and alarming indicator of severe conditions, including dementia, stroke, and brain tumors (American Psychiatric Association, 2022; Dara et al., 2014; Dara et al., 2013; Durfee et al., 2021; Leyton & Hillis, 2017; Scherer, 2003; Wright et al., 2018).

### The Nature of Linguistic Prosody

If, as generally assumed, linguistic prosody is both functionally and structurally different from affective prosody (Caplan, 1987b), it should display distinct structural and functional characteristics concerning the corresponding ROI activations. More specifically, and the primary motivation for distinguishing linguistic prosody from affective prosody, is the role it plays in language and the distinct predictions that are pre-supposed by this. As part of the linguistic system, prosody is language-specific rather than universal. These led to works on prosodic typology (Jun, 2005; Jun, 2014) and a plethora of specialized works on the prosody and intonation of various languages, such as English (Liberman, 1979; J. Pierrehumbert, 1980), Catalan (Prieto, 2009), Swedish (Bruce, 1977), Italian (D’Imperio, 2001; D’Imperio et al., 2014), Greek (Arvaniti et al., 1998; Themistocleous, 2016b), and Japanese (Beckman & Pierrehumbert, 1986b). Also, linguistic prosody, unlike affective prosody, should provide a certain type of domain specificity, manifested by hierarchical and syntagmatic relationships—typical for language systems like morphology and syntax—that determine prosodic patterns in utterances, namely the prosodic units and “rules” or patterns that explain how the prosodic units are ordered and structured hierarchically. Prosodic units are categorical as they are part of the language system.

Given the systematic redundancy and selectivity, which characterizes the linguistic systems, certain prosodic functions can be performed by other linguistic domains of grammar and vice-versa. For example, *contrast* in English is expressed through prosody “GEORGE did not go the cinema; MARY did” or through a combination of syntactic and prosodic forms, such as cleft sentences “It was GEORGE who went to the cinema (not Anna).” The linguistic selectivity also accounts for the significant cross-linguistic variation of prosodic functions. For example, in English new information focus and topics (known information) manifest with tonal movements, such as nuclear pitch accents for focus or prenuclear pitch accents for topics whereas Korean and Japanese employ morphological topic markers to designate the constituents in topic (Jun, 2015; Themistocleous, 2011).

Linguistic prosody covers both the lexical prosody and post-lexical prosody, although it is important to distinguish between both concerning their function and role in language.

### Lexical Prosody

Lexical prosody associates with word-level distinctions, such as stress in English, as in words like ínsert and insért, pitch accents in Norwegian bønner /bœnnər/ (beans) and bønder /bœnnər/ (farmers), and tones in Mandarine Chinese 妈 mā mother, 麻 má hemp, 马 mǎ horse, 骂 mà scold. Lexical prominence can distinguish word meanings, as English or Greek, or has a fixed position in the word in languages such as Czech and Hungarian, in this case its role is to differentiate each word from another. Trubetzkoy (1969) distinguished tonal phenomena into cumulative functions (indicating prominence) and delimitative functions (marking domain boundaries) (Beckman, 1986)^2^. There has been a strong debate in the literature on the acoustic manifestation of lexical prominence and its acoustic and perceptual correlates. For example, earlier research suggested the fundamental frequency (*F0*) is the acoustic correlate of intonation ^3^ and duration of quantity (Stevens, 1998). Nevertheless, the dynamic nature of speech production involves an interplay between several acoustic properties, such as an increase of syllable duration, intensity, and a certain *F0* movement—e.g., a rise or a fall of the fundamental frequency— marking anchored on the stressed syllable. In pitch accent languages, such as Japanese and Norwegian, syllable length is phonemic, namely it distinguishes long and short syllables. For example, in Norwegian there are minimal pairs of long and short vowels is denoted orthographically: words that end in double consonant are preceded by a short vowel whereas those with a singleton by a long vowel, e.g., pen /ˈpeːn/ ‘pretty’ vs. penn /ˈpɛn/ ‘pen’, fet /ˈfeːt/ ‘oily’ vs. fett /ˈfɛtː/ ‘fat’, and blek /blˈeːk/ ‘pale’ vs. blekk /blˈɛkː/ ‘ink’.

### Postlexical Prosody

The postlexical prosody has three primary functions: i) it highlights post-lexical constituents (e.g., words, phrases or utterances), ii) demarcates their boundaries, and iii) distinguishes the melodies of utterances, (e.g., statements, questions, and commands). For example, by using different pitch accents, the speaker of Greek in Figure 1 modifies the type and placement of emphasis (a.k.a., focus) in the first word (A) and the second word (B) in polar questions. The Greek language distinguishes polar questions and statements through intonation primarily.

**Figure 1.**
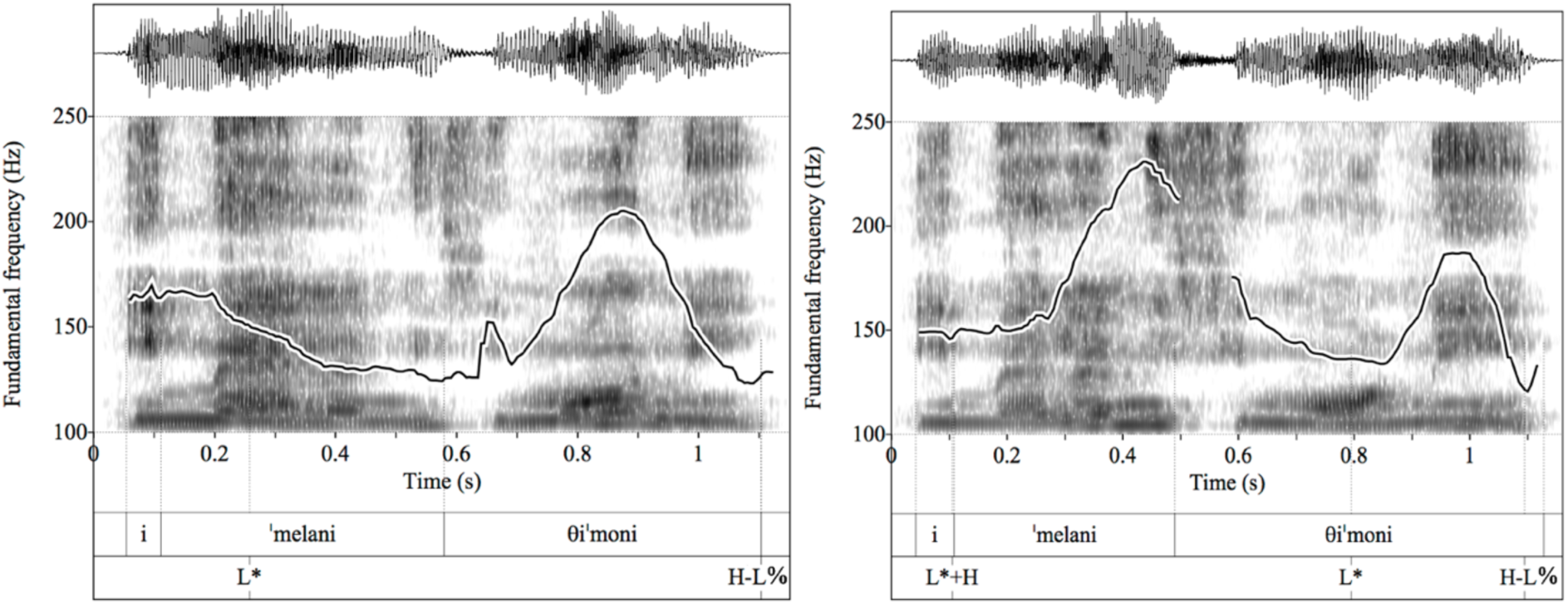
The image shows the waveform (top tier), the spectrogram with a superimposed fundamental frequency contour (F0) in Hertz (Hz) (middle tier), the segmental and prosodic annotation in ToBi for two different sentences, η Μέλανη θυμώνει “Melanie gets angry”. Τhe first polar (yes or no) question is uttered with a focus on the first word, and the second with a focus on the last word. Greek polar questions are only marked prosodically.

We should expect that speakers organize post-lexical prosodic constituents in hierarchical prosodic domains (Figure 2). In oral speech, speakers may break their utterances in ways that do not correspond to the syntactic constituents, such as phrases and sentences and form separate prosodic unities (Shattuck-Hufnagel & Turk, 1996). There is a consensus over the following: syllables, feet, prosodic words, intermediate phrases, intonational phrases (Beckman & Pierrehumbert, 1986a; Hayes, 1995; Nespor & Vogel, 2007; Selkirk, 1984, 1995; Shattuck-Hufnagel & Turk, 1996; Wagner, 2005). A foot is a rhythmic unit, composed of stressed and unstressed syllables. A prosodic word is a word or a combination of words with a single primary stress. One or more intermediate phrases typically conclude with a rising intonation, creating an intonational phrase—a larger unit of speech melody. The domains are delimited by various phenomena.

**Figure 2.**
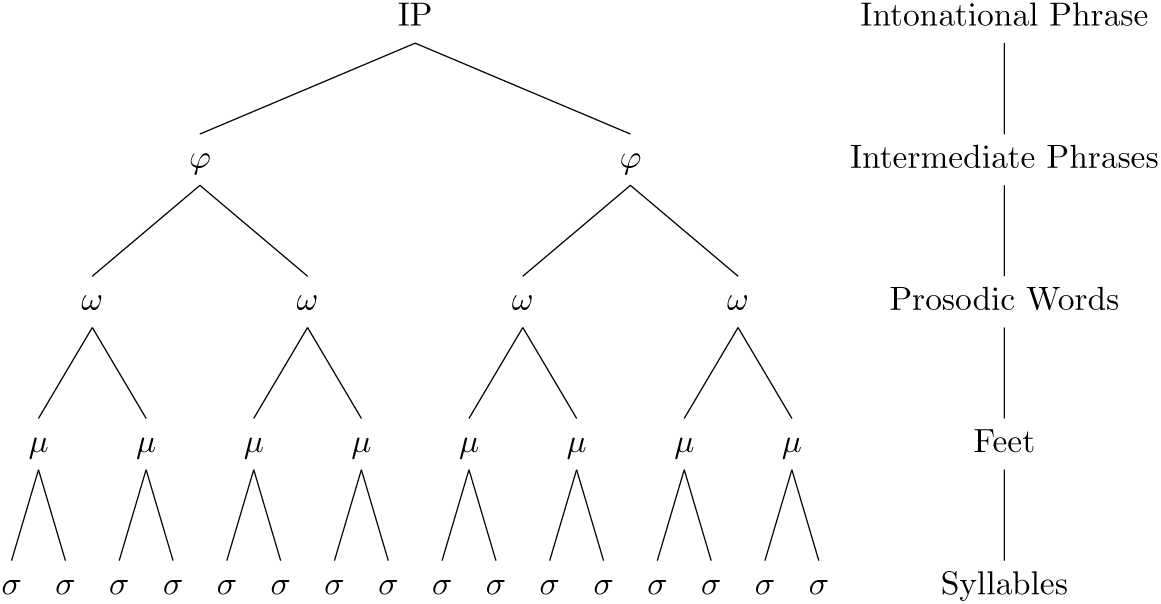
Abstract prosodic hierarchy, shown matches of phonological/prosodic domains from syllable to intonational phrases. Abstract prosodic hierarchy, shown matches of phonological/prosodic domains from syllable to intonational phrases.

The Intermediate and Intonational Phrases are high-order domains, superseding the prosodic word. Each of these two domains carries a main tonal prominence and a delimitative tonal movement demarcating primarily the corresponding right boundary. The tonal prominence is manifested by the pitch accents, which associate with the stressed syllables. Traditionally, the pitch accent that conveys the primary prominence or focus of the intermediate or intonational phrase is *the nuclear pitch accent* whereas the others are the prenuclear pitch accents. ^4^ The prenuclear pitch accents are commonly associated with the known or old information in an utterance (topic) whereas nuclear pitch accents are associated with the new information focus or contrastive information on different constituents such as syllables, morphemes, words, phrases ([F], indicates the focused materials) (Themistocleous, 2016b). (1)

- I said PRE[F]cede | not PRO[F]ced
- I[F] went to the cinema | not YOU[F]!
- I went to the CINEMA[F] | not to the MALL[F]!
- John bought a NEW[F] car. “New information about the car, e.g., Did John buy an old car?”
- John BOUGHT[F] a new car. “Highlighting the act of buying, What did John do?”

JOHN[F] bought a new car. (Highlighting the subject, John, e.g., Who bought a new car?)

In addition to the pitch accents, the phrase accents and boundary tones mark the right edges of Intermediate and Intonational Phrase correspondingly. A critical insight by Beckman and Pierrehumbert (1986a, 1986b) was that the right edge of Intonational Phrases is not marked by one tonal movement but two, namely, they suggested that the right edge patterns can be explained by a phrase accent, which marks the boundaries of the intermediate phrase and a boundary tone marking the boundaries of the intonational phrase.

Overall, there is a considerable variability of prosodic phenomena that delimitate the boundaries of higher-order domains, such as the intonational and intermediate phrases. These delimitative phenomena include rises or falls of the F0, final lengthening, pauses, and a lowering of the intensity (Themistocleous, 2014). Phenomena, often with social identity signaling (Yuasa, 2010), such as changes in voice quality, aspiration, and nasalization often delimitate prosodic domains (Fox, 2000; Lehiste, 1970). For example, a low creaky voice (vocal fry) with lowering of F0 at the right edge of the utterance, is more common among young women. These salient phenomena depend on the underlying cognitive phonological representation and on the mechanics of articulation (e.g., airflow, pressure, articulatory speed, and slowing down at the end of speech (Beckman & Edwards, 1990; Byrd & Saltzman, 1998; Cooper & Paccia-Cooper, 1980; de Jong et al., 1993; Fougeron & Keating, 1997; Turk & Shattuck-Hufnagel, 2000). Finally, phonological phenomena such as such as assimiliation, are being blocked within prosody word boundaries the e.g., [n] > [m] is block post-lexically, in “Who’s in? Possibly Mary and George” /huːz **ɪn ˈp**ɒsəbli ˈmɛəri ænd ˈdʒɔrdʒ/ (but [n]>[m] “impossible” /ɪ**mˈp**ɑsɪbəl/ “in” + “possibly”).

### A Linguistic Prosody Representation System

A development of the research on prosody in the 1970 and 1980s (Bruce, 1977; Liberman, 1975; J. B. Pierrehumbert, 1980) was the specification of tonal phonology and the establishment of the tone and boundary indices transcription system (ToBI). The basic idea is that the rises and falls of the melodic patterns involve two abstract tonal targets: a high (H) and a low (L) tone^5^. *Pitch accents* consist of an L or H, or a combination of the two: L*, H*, L+H*, L*+H, H+!H*.^6^ Common pitch accents in English ToBI include:

- *H* (high):* A high pitch target on the stressed syllable.
- *L* (low):* A low pitch target on the stressed syllable.
- *L-H* (low-high):* A rise from a low to a high pitch on the stressed syllable.
- *H-H* (high-high):* A rise from a high to a low pitch on the stressed syllable.

The star symbol (*) designates the tone that is timed with the stressed syllable (e.g., H*L) (Beckman et al., 2005). Phrase accents are indicated as high (H-), downstepped high (!H-), and low (L-) mark the right edge of intermediate phrases. The boundary tones, high (H%), low (L%), mark the end of the phrase accents. Thus, the end of intonational phrases is demarcated by a combination of a phrase accent marking the end of the intermediate phrase and a boundary tone that marks the right edge of the intonational phrases (when there is only one intonational phrase, that coincides with the intermediate phrase). Thus, there can be several possibilities at the end of an intonational phrase:

- L-L% A low boundary tone marking the right edge boundary.
- L-H% (*a low phrase and a high boundary tone*): a rising melodic pattern at the end of the intonational phrase

As the *F0* declines over the course of the utterance, due to a drop in the energy and sub-laryngeal pressure, a high pitch accent (H*) at the utterance’s right edge can be lower than a preceding L* accent, which is compensated during speech perception.

One additional aspect of ToBI is that it denotes phenomena at the boundaries of prosodic domains through the Boundary Indices from 1 to 5, which indicate how strong is the connection between those domains.

### The Nature of Affective Prosody

The emotional aspects of prosody received less attention from both linguists, who are more likely to consider affective prosody as non-linguistic or paralinguistic (Ladd, 2009) and aphasia researchers, who classify affective prosody among the non-core or peripheral aspects of language functioning (Caplan, 1987a, p. 389). Furthermore, affective prosody is not well-described within the linguistic theory. It is often employed in the literature to include diverse speech communication phenomena, such as emotions, it (e.g., expressions of joy and anger), prosodic styles of speech (e.g., high pitched speech often used for politeness and irony), turn-taking cues in conversations (Sacks et al., 1974), and social indexical connotations, the subtle prosodic cues conveying social information (Eckert, 2019; Foulkes & Docherty, 2006; Silverstein, 2010).

Despite the limitations, there has been a lot of research on the affective aspects of prosody concerning prosody impairment. Dysprosody and aprosodia where first discussed by the Norwegian neurologist Monrad-Krohn (1947), who coined the two terms to refer to impaired prosody (dysprosody) and the latter lack of prosody (aprosodia). Expanding on that foundational work, Ross (1981) identified two primary types of aprosodias, receptive aprosodia “difficulty to perceive prosody” and expressive aprosodia “difficulty to express emotions”. Individuals with receptive aprosodia have difficulty understanding the emotional content conveyed through prosody, such as identifying whether an utterance is spoken in a happy, sad, or angry tone. Individuals with expressive aprosodia have difficulty conveying emotions through their own speech. Their speech may sound monotone or flat (Ross, 1981), even when they are trying to express emotions. Common related prosodic subtypes include *sensory aprosodia*, which refers to the impaired affective prosody recognition and imitation, but spared prosodic production in spontaneous speech; *transcortical sensory aprosodia*, which refers to the impaired affective prosody recognition, but spared repetition and spontaneous production; *motor aprosodia*, which is the impaired affective prosody production in spontaneous speech and repetition, without impaired comprehension; and the *transcortical motor aprosodia*, where there is intact repetition and comprehension, but poor spontaneous use of affective prosody. Other subtypes, including *intellectual* and *assertive prosody, which as stated above denote* prosodic styles of speech rather than emotions (Ross & Mesulam, 1979). Individuals with ischemic brain damage due to stroke and Multiple Sclerosis (MS), often manifest aprosodias, which are “disorders of affective language that occur following right-hemisphere damage” (Ross, 1981). Impairments, such as monotonous speech, can be indicative of neurological conditions like depression, Alzheimer’s disease, Parkinson’s disease, Primary Progressive Aphasia, and other types of dementia (Jonathan D. Rohrer et al., 2012).

### Brain Areas involved in Prosody

Linguistic prosody is part of the language network and gives meaning to the continuous range of frequencies perceived from the environment. Basic cognitive processes are initiated before the activation of the language networks, such as the pitch discrimination, the temporal properties of frequencies, and the direction of tonal changes are typically established in the acoustic cortex and the frontal areas of the brain (Angenstein et al., 2012; Johnsrude & Rodd, 2016). These processes follow the general predictions of speech perception, and production are modeled by the dual stream model of speech perception (Hickok & Poeppel, 2007). The model involves a cortical organization of speech within two main streams of information: the dorsal and the ventral stream (Hickok & Poeppel, 2007). The dorsal stream facilitates speech production by mapping the language structures to articulation. The ventral stream facilitates speech perception and is responsible for associating a meaning to sounds. The ventral stream initiates bilaterally, which explains activations of linguistic prosody in both hemispheres, although most of the linguistic processed are eventually active in the left hemisphere (Hickok & Poeppel, 2007). On the other hand, affective prosody is localized in the right hemisphere. In patients with right hemisphere stroke, affective prosody impairment depends on the type and timing of stroke. Studies in acute and chronic stroke phases are valuable for understanding the neural mechanisms of affective prosody. Acute studies can help identify the initial impact of the lesion, while chronic studies can shed light on the long-term consequences and potential for recovery. For example, studies such as Durfee et al. (2021) and other studies, as we will see in the subsequent section, associate impaired affective prosody with a RH stroke, which suggests that there is a strong involvement of RH areas in the perception and production of affective prosody.

### Previous Reviews

Several recent narrative reviews have addressed the neural substrates of prosody (Kemmerer, 2014, pp. 189-212; Kotz et al., 2006; Pell, 2006; Schirmer & Kotz, 2006; Wildgruber et al., 2006; Wildgruber et al., 2009). First, the hemispheric specialization and expression of prosody has been explored by earlier meta-analysis reviews. Ukaegbe et al. (2022) focused on studies of aprosodia following focal left- or right-hemisphere damage, showing inconsistent support for selective contribution of the two cerebral hemispheres during comprehension and production of prosody. However, their findings provided strong evidence that RH lesions have the potential to disrupt emotional prosody comprehension to a greater degree than left hemisphere (LH) lesions (Ukaegbe et al., 2022). With respect to the neural substrates of linguistic and affective prosody concerning neural pathways and networks the answers are again inconclusive. A meta-analysis by Witteman et al. (2012) argue that there is not a shared conclusion from the studies, they reviewed. Moreover, Witteman et al. (2012) pointed to three main limitations of their meta-analysis, the limited number of participants in the reviewed papers, and two methodological constraints of the review, the limited numerical of participants and that they could not consider the effect sizes or the extend of activation but only the peak-coordinates of activation clusters. Belyk and Brown (2014) found brain activation in the right-hemisphere auditory areas. BA 47 is more likely activated for affective prosody productions, while linguistic prosody tends to trigger activations in the ventral part of BA 44. A more recent systematic review by Durfee et al. (2021) evaluated the localization of brain lesions that impair affective prosody, which is the changes in pitch, rhythm, and loudness of speech that convey emotion. They found that damage to right antero-superior (dorsal stream) regions of the brain was associated with impairments in affective prosody production, while damage to more posterolateral (ventral stream) regions resulted in affective prosody comprehension deficits. The authors concluded that distinct RH regions are vital for affective prosody comprehension and production.

## The Current Review

### Scope and Aims of the Review

Given the strong theoretical and practical implications of determining the cortical substrates of prosody, we decided to conduct a systematic meta-analytic review of existing studies. We assess the extent to which linguistic and affective prosody differ in the corresponding cortical areas that indicate stronger associations and examine especially the areas that might be unique in these two cases, which can indicate their different manifestation.

## Method

We will use an ALE meta-analysis to detect robust activations in papers reporting affective and linguistic prosody and answer the research questions in the paper. Papers were selected for the meta-analysis according to the PRISMA (Preferred Reporting Items for Systematic Reviews and Meta-Analyses) guidelines and the PRISMA 2020 checklist, available at www.prisma-statement.org (Page et al., 2021). PRISMA provides a structured approach based on consensus for developing systematic reviews and meta-analyses. Subsequently, the ALE meta-analysis was conducted (Turkeltaub et al., 2002).

### Literature Search, Inclusion Criteria, and Coder Reliability

In this meta-analysis, we searched Embase, NeuroQuery. Oria, PsycNet, PubMed (NLM), Scopus, and Web of Science until 2024 July 20. Our search criteria did not specify the language of the studies published but we included only works in languages that we can safely review, namely Danish, English, French, Greek, Italian, and Norwegian, Swedish. When locating studies, we used combinations of keywords related to prosody (*“prosody”, “intonation”, “pitch accent”*), prosody impairment (“aprosodia” OR “dysprosody”) crossed with terms related to the brain (“brain”, “cortex”, “subcortical structures”) and neuroimaging/neurophysiology techniques (“MRI” OR “EEG” OR “EcOG”). The data were imported into EndNote 21 (The EndNote Team, 2013), which facilitated the selection of the final papers for review (Figure 3).

**Figure 3.**
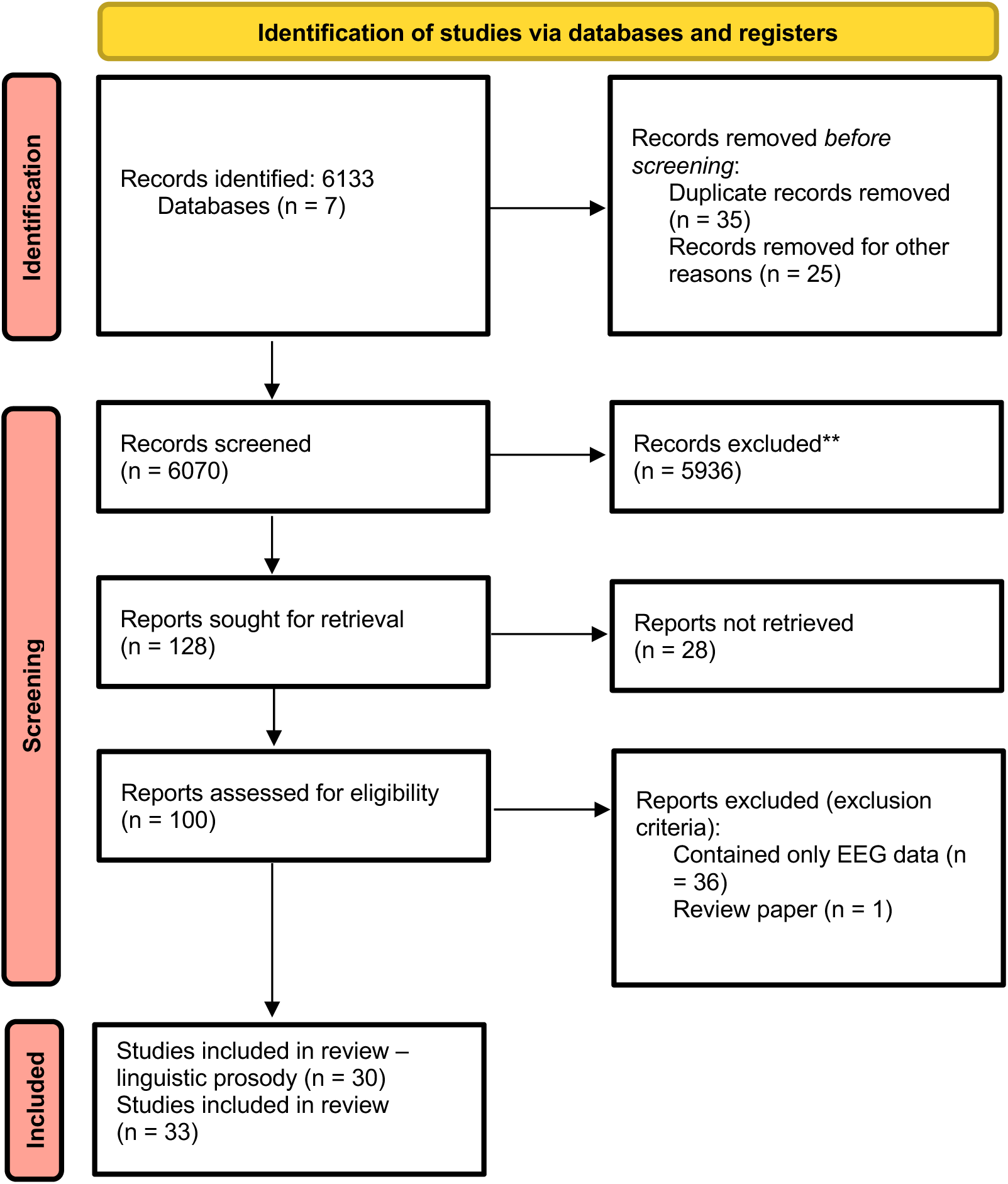
Prisma diagram showing the records identified, screened, and included in this study (Page et al., 2021).

The selection of the studies followed a comprehensive screening of the collected articles based on the inclusion and exclusion criteria for the review and analysis (Table 2). Although we researched primarily peer-reviewed literature, we analyzed publication results from the gray sources as well, including the PhD theses and non-peer-reviewed conferences. These were not included in the review but provided an additional perspective on the studies and the findings in the area.

**Table 1.** Database Search Query.

| Search term combinations for PubMed (NLM) |
| --- |
| ((prosody OR intonation OR suprasegmentals OR pitch accent OR aprosodia OR dysprosody) AND (cortex OR brain OR subcortical structures) AND (MRI OR EEG OR EcOG)) |

**Table 2.**
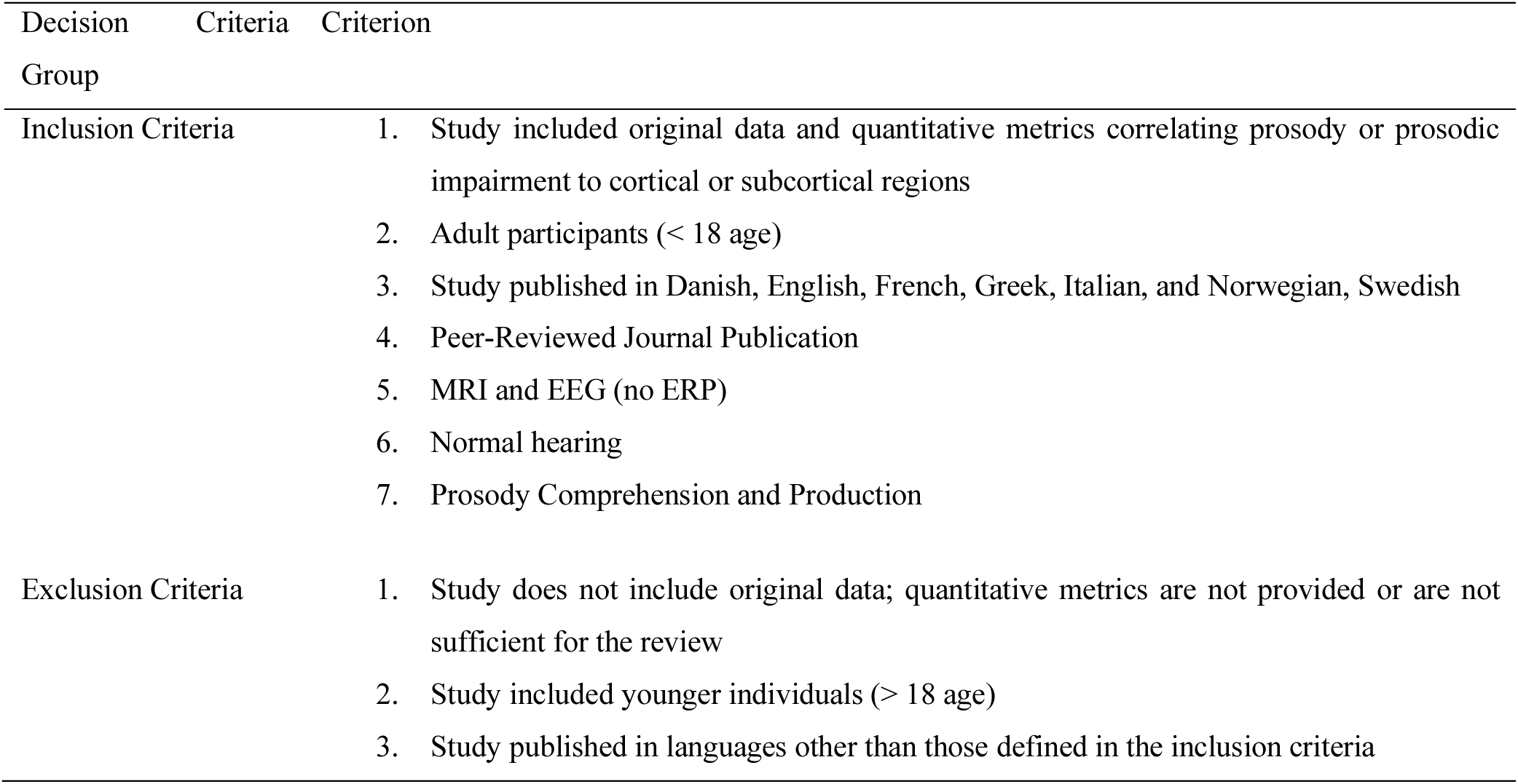
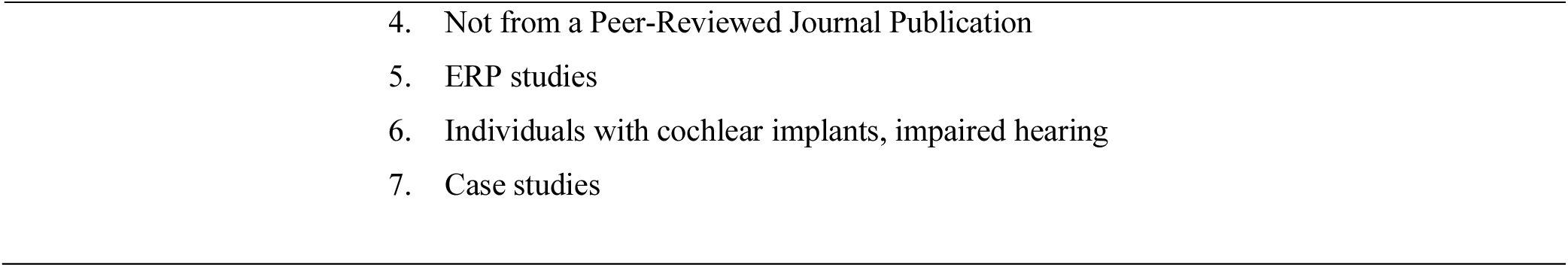
Inclusion and Exclusion Criteria.

We have selected the reported MNI space both from the tables and the text and employed those for the ALE meta-analysis.

### ALE meta-analysis

Using the ALE approach to systematic meta-analysis, we evaluate the MRI activations reported in studies on affective and linguistic prosody to provide a new representation of the corresponding effectiveness of the combined results (Turkeltaub et al., 2002). Each activation focus reported by one study is represented as a 3D Gaussian probability distribution instead of a single point. The number of participants adjusts the spread of the Gaussian distribution, and subsequently, an “activation likelihood map” for each study is generated. The smaller studies have a wider distribution, whereas studies with more participants have a narrower distribution. Narrower distributions indicate more concentration of values towards the center of the distribution and thus are more confident that the reported activations are part of the distribution. For every voxel in a study, the “activation likelihood map” provides the highest probability of activation from the modeled activation foci. Turkeltaub et al. (2002), introducing the ALE meta-analysis, employed a Random-Effects Analysis, evaluated the convergence of activation likelihood points in the studies included in the paper, and detected those points that were greater than what would be expected by chance if activations were randomly distributed. Subsequently, Turkeltaub et al. (2002) provided a statistical map showing the regions where activations converged; in other words, the regions where activations overlap are significantly greater than chance. Therefore, the generated new map includes activations that were consistent across papers. Recent developments in the ALE meta-analysis are more sensitive to the effects of the number of participants and the limitations of small sample sizes.

The specific framework of ALE-meta-analysis employed in this paper is based on the standard protocol implemented using the brainmap set of tools, primarily the GingerALE tool (Turkeltaub et al., 2012), which employs an updated version of the ALE based on Eickhoff et al. (2009). Specifically, the GingerALE first calculates the ALE score for each voxel and assesses the null distribution of each voxel’s ALE score. Then, it estimates the threshold for the ALE image using the *p*-values. We had selected a Voxel-based Family-wise Error permutation set to 0.01, a permutation threshold selected to 1000, and a minimum volume set to 200 mm^3^. Clusters were found from a cluster analysis on the thresholded map, according to the GingerALE protocol, and associated with the corresponding Talairach label for the area, which is reported in Tables 3 to 5. We employed FSLeyes 1.12.3 to overlay the GingerALE on the MNI152_T1_1mm standard (McCarthy, 2024).

## Results

We present the results on the cortical correlates of linguistic prosody first. Subsequently, we report on the outcomes for affective prosody and the contrasts between linguistic and affective prosody. The detailed selection and characteristics of the included are reported in Appendix 1 for linguistic prosody and Appendix 2 for affective prosody.

### Cortical Correlates of Linguistic Prosody

Figure 4 and Table 3 present the findings from a Ginger ALE meta-analysis on brain regions associated with linguistic prosody. The analysis identifies several clusters of brain activation associated with linguistic prosody, involving both the frontal and temporal lobes, with most regions being in the right hemisphere. Specifically, the key Brain Regions involved are the following

1. *Right Frontal Lobe. i.* Medial Frontal Gyrus (BA 8): Cluster 1 shows a peak activation at coordinates (8, 24, 42) with an ALE value of 0.035. This region is implicated in executive functions and decision-making. ii. Superior Frontal Gyrus (BA 6): Another peak within Cluster 1 is located at (10, 14, 50) with an ALE value of 0.033, suggesting involvement in motor planning and control.
2. *Left Superior Temporal Gyrus.* This Cluster 2 contains two peaks. One at coordinates (-54, -4, -2) with an ALE value of 0.039, indicating significant activation in the superior temporal gyrus, a region involved in auditory processing. Another peak at (-58, -14, 2) with an ALE value of 0.028, also within the superior temporal gyrus, associated with Brodmann area 22, which is linked to language comprehension.
3. *Right Frontal Lobe.* Precentral Gyrus (BA 6): Cluster 3 shows a prominent peak at (54, 2, 42) with the highest ALE value of 0.042, indicating strong activation in the precentral gyrus, a region involved in motor control.
4. *Right Sub-lobar Region.* Insula (BA 13): Cluster 4 shows a peak at (34, 22, 4) with an ALE value of 0.033, suggesting involvement of the insula, a region implicated in emotional and sensory integration. All clusters exhibit highly significant ALE scores (ranging from 0.028 to 0.042) with p-values of 0.000, indicating robust evidence of brain activation associated with linguistic prosody across these regions.

**Figure 4.**
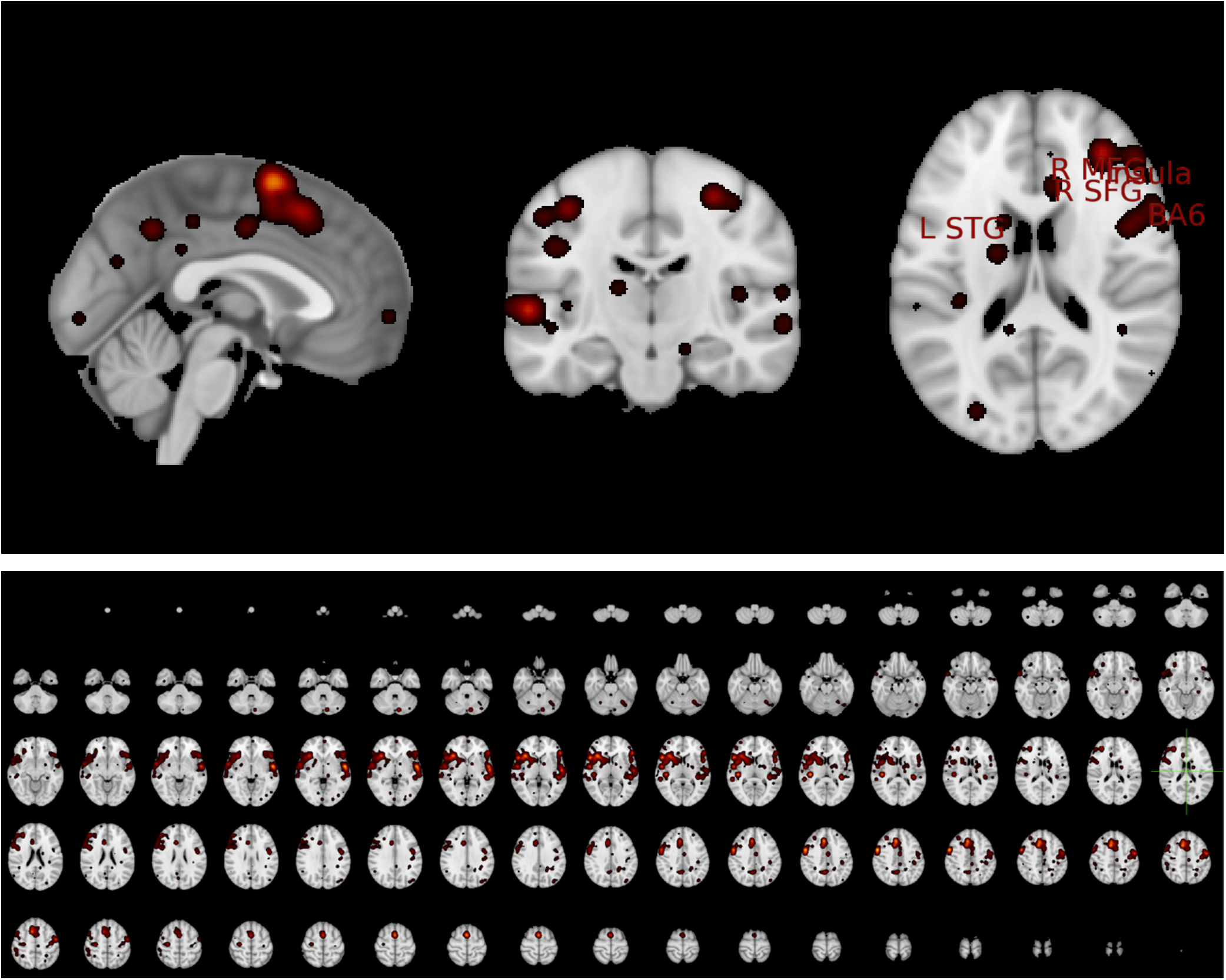
Orthographic view showing the sagittal, coronal, and axial views of the activated areas in the MNI space from the ALE meta-analysis. The labels indicate activations of linguistic prosody provided by the cluster analysis of the ALE meta-analysis (Top Panel) and the lightbox view (Bottom Panel). The view has been centered using standard settings at x= -0.5, y -17.5, z=18.5.

The identified brain regions suggest that linguistic prosody engages both motor and auditory processing areas, along with regions involved in emotional and sensory integration.

### Affective Prosody

The results of a Ginger ALE meta-analysis on the relationship between affective prosody and brain activation are summarized in Figure 5 and Table 4. The analysis identified six clusters of brain regions associated with affective prosody. These clusters are primarily located in the superior temporal gyrus (both left and right), inferior frontal gyrus, and the parahippocampal gyrus near the amygdala.

1. Two clusters (-62, -24, 4) and (-58, -18, -2) indicate strong activations in with ALE values of 0.055 and 0.048, respectively related to the *Left Superior Temporal Gyrus (BA 22)*.
2. A single cluster (-50, 26, 6) with an ALE value of 0.059 represents a peak activation in the Left Inferior Frontal Gyrus (BA 45).
3. Peaks in clusters 3 (58, -6, -6 and 58, -12, -2) and cluster 4 (64, -26, 2), with ALE values ranging from 0.048 to 0.054, indicating consistent activation in the *Right Superior Temporal Gyrus (BA 22)*.
4. A notable peak in cluster 5 (-20, -6, -14) with a high ALE value of 0.059, suggesting a strong association with affective prosody, is an activation in the *Left Parahippocampal Gyrus – Amygdala*.
5. Two peaks in cluster 6 (48, -32, 4 and 48, -26, 4) with lower ALE values (0.045 and 0.044), indicating moderate activation in the *Right Superior Temporal Gyrus (BA 41)*.

**Figure 5.**
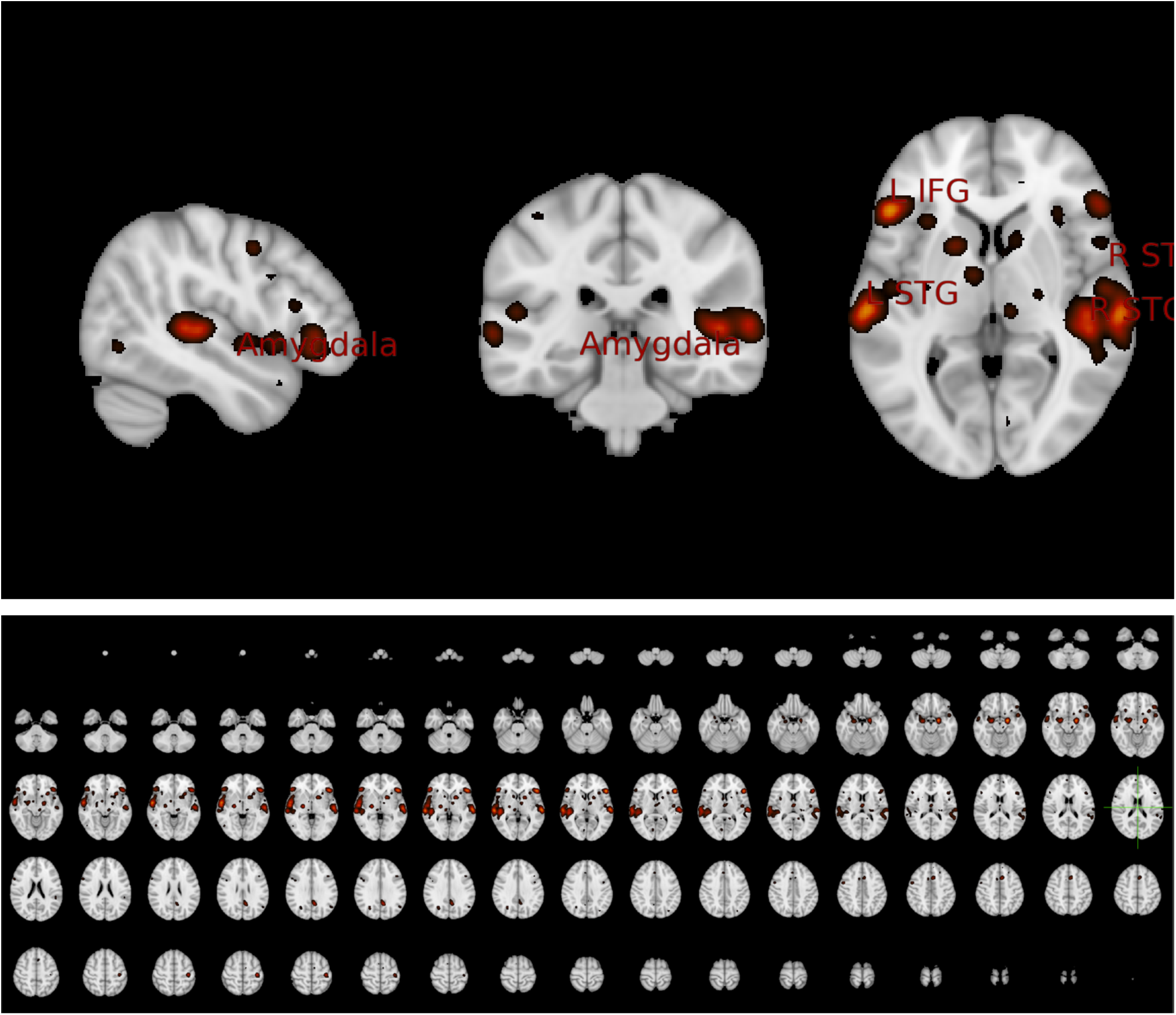
Orthographic view showing the sagittal, coronal, and axial views of the activated areas in the MNI space from the ALE meta-analysis. The labels indicate activations of affective prosody provided by the cluster analysis of the ALE meta-analysis (Top Panel) and the lightbox view (Bottom Panel). The view has been centered using standard settings at x= -0.5, y -17.5, z=18.5.

**Table 3.** Cluster analysis form the ALE meta-analysis and peak outcomes for linguistic prosody.

| Cluster # | x | y | z | ALE | P | Z | Label (Nearest Gray Matter within 5mm) |
| --- | --- | --- | --- | --- | --- | --- | --- |
| 1 | 8 | 24 | 42 | 0.035 | 0.000 | 6.407 | Right MFG BA8 |
| 1 | 10 | 14 | 50 | 0.033 | 0.000 | 6.139 | Right SFG BA 6 |
| 2 | -54 | -4 | -2 | 0.039 | 0.000 | 6.907 | Left STG |
| 2 | -58 | -14 | 2 | 0.028 | 0.000 | 5.538 | Left STG BA 22 |
| 3 | 54 | 2 | 42 | 0.042 | 0.000 | 7.213 | Right PCG BA 6 |
| 4 | 34 | 22 | 4 | 0.033 | 0.000 | 6.177 | Right Insula BA 13 |

SFG: Superior Frontal Gyrus; MFG: Medial Frontal Gyrus; STG: Superior Temporal Gyrus; PCG: Precentral Gyrus.

**Table 4.** Cluster analysis form the ALE meta-analysis and peak outcomes for affective prosody.

| Cluster # | x | y | z | ALE | P | Z | Label (Nearest Gray Matter within 5mm) |
| --- | --- | --- | --- | --- | --- | --- | --- |
| 1 | -62 | -24 | 4 | 0.055 | 0.000 | 7.211 | Left STG (BA 22) |
| 1 | -58 | -18 | -2 | 0.048 | 0.000 | 6.524 | Left STG (BA 22) |
| 2 | -50 | 26 | 6 | 0.059 | 0.000 | 7.598 | Left IFG (BA 45) |
| 3 | 58 | -6 | -6 | 0.054 | 0.000 | 7.131 | Right STG (BA 22) |
| 3 | 58 | -12 | -2 | 0.048 | 0.000 | 6.538 | Right STG (BA 22) |
| 4 | 64 | -26 | 2 | 0.054 | 0.000 | 7.137 | Right STG (BA 22) |
| 5 | -20 | -6 | -14 | 0.059 | 0.000 | 7.555 | Left PHG - Amygdala |
| 6 | 48 | -32 | 4 | 0.045 | 0.000 | 6.216 | Right STG (BA 41) |
| 6 | 48 | -26 | 4 | 0.044 | 0.000 | 6.146 | Right STG (BA 41) |

Notes: IFG: Inferior Frontal Gyrus; SFG: Superior Frontal Gyrus; MFG: Medial Frontal Gyrus; STG: Superior Temporal Gyrus; PHG: Parahippocampal Gyrus.

All clusters showed highly significant ALE scores (ranging from 0.044 to 0.059) and corresponding p-values of 0.000, indicating robust findings across these regions. The nearest gray matter locations within 5mm of the identified coordinates suggest that affective prosody involves regions typically associated with auditory and language processing (superior temporal gyrus), as well as emotional processing (parahippocampal gyrus - amygdala).

### Contrasts

We found contrasts only concerning linguistic – affective prosody. Figure 6 and Table 5 show the results on the contrast linguistic – affective prosody.

**Figure 6.**
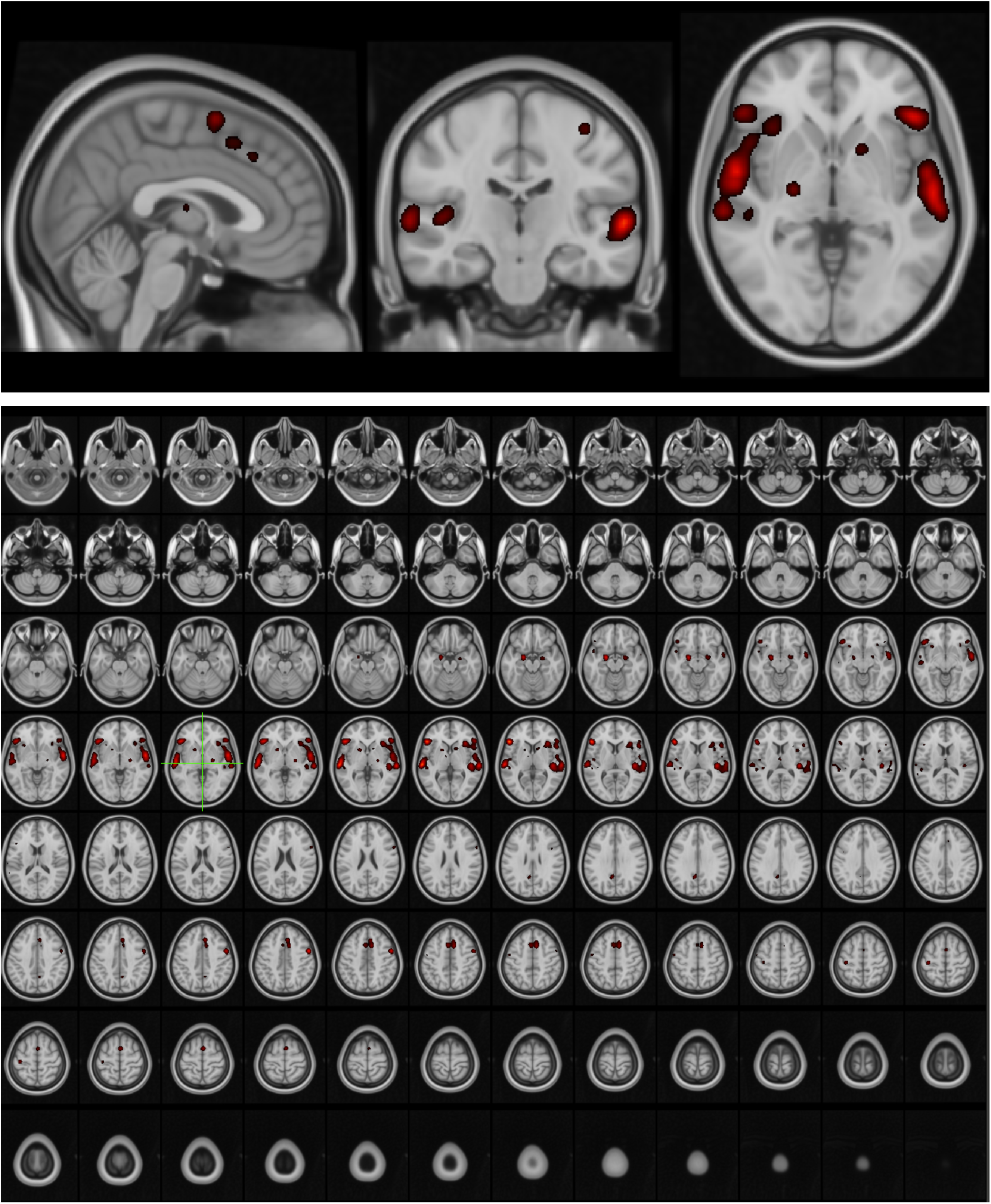
Orthographic view showing the sagittal, coronal, and axial views of the activated areas in the MNI space from the ALE meta-analysis for the contrast linguistic – affective prosody. The labels indicate activations of affective prosody provided by the cluster analysis of the ALE meta-analysis (Top Panel) and the lightbox view (Bottom Panel). The view has been centered using standard settings at x= -0.5, y - 17.5, z=18.5.

**Table 5.** Cluster analysis form the ALE meta-analysis and peak outcomes for the contrast linguistic – affective prosody.

| Cluster # | x | y | z | P | Z | Label (Nearest Gray Matter within 5mm) |
| --- | --- | --- | --- | --- | --- | --- |
| 1 | 6 | 20 | 44 | < 0 | 3.29 | Right Medial Frontal Gyrus BA |

## Discussion

Prosody refers to the melodic patterns of speech involving pitch, loudness, quantity, speech rate, and voice quality, including features such as aspiration and nasalization. Clinical research distinguished between affective (emotional) and linguistic prosody. Affective prosody expresses meanings like anger, joy, and sadness, and it is relatively consistent across cultures. The linguistic prosody, which is part of the language system (Caplan, 1987a) is further subdivided into lexical prosody, which involves language-specific prominences: word-level stress, tone, and pitch accents, and post-lexical prosody, which operates across phrases and sentences, using intonation to highlight meanings, express utterance tunes, and organize the utterance into units.

This ALE meta-analysis study investigated the cortical activations of affective and linguistic prosody reported in the literature on healthy individuals to identify the cortical correlates and activation patterns related to linguistic and affective prosody. The findings provided strong supporting evidence about the neural correlates of affective and linguistic prosody and clarified their relationship. Also, it showed a strong bilateral activation related to linguistic and affective prosody. We believe that the relational findings, in turn, have important theoretical implications on the role of prosody in speech perception and practical impacts on designing prosodic treatment approaches.

Specifically, in the studies we have reviewed, linguistic processing (e.g., phonology, morphology, syntax, and semantics), including lexical prosody and the integration of prosody to language, are strongly related to left-lateralized brain areas (van der Burght et al., 2019; van Leeuwen et al., 2014) whereas right-lateralized areas are involved in processing emotional prosody (Ethofer et al., 2009; Sammler et al., 2015; Tracy et al., 2011). For example, several studies have reported activation of both hemispheres and argue that the right hemisphere is more sensitive to prosodic variations, while the left hemisphere is employed in processing linguistic aspects of pitch (Inspector et al., 2014; Kandylaki et al., 2017; Meyer et al., 2004). The left IFG is responsible for processing intonational cues linked to language during sentence comprehension, suggesting a left hemisphere dominance in the language (van der Burght et al., 2019). Also, the left IFG is responsible for tonal processing during sentence comprehension, guiding sentence processing (Chien et al., 2021; van der Burght et al., 2019). Moreover, van Leeuwen et al. (2014) suggest that the left IFG sequences pitch accents that signal information structure during sentence comprehension. The left IFG and pSTS are contributing to the interaction of prosody in organizing syntactic domains and sequencing, especially in relation to the early-closure garden-path syntactic processing and the reanalysis of sentences with prosodic and plausibility conflicts (den Ouden et al., 2016).

However, not all studies agree on this clear-cut specification of the two hemispheres. Other studies show the contribution of both hemispheres in both affective and linguistic prosody (Inspector et al., 2014; Kandylaki et al., 2017). Importantly, as shown in Table 6, the prosody related activations, derived from this metanalysis, manifest bilaterally in both the linguistic and affective prosody and after evaluating the contrasts, we found only one peak activation that was different the Right Medial Frontal Gyrus. One account for these observations is that a bilateral involvement is due to early activation of both left and right frontotemporal areas during speech perception (Hickok, 2013; Hickok & Poeppel, 2007). Specifically, high and low frequencies create distinct responses in the inner ear. High and low frequencies cause deformations of the cochlear hair cells, allowing them to respond to different pitch frequencies from high to low: high frequencies are represented tonotopically at the base of the basilar membrane and lower frequencies at the interior and closer to the apex. As predicted by the dual stream model for speech perception, a spatiotemporal reception of the sound occurs in the left and right superior temporal gyrus (Hickok & Poeppel, 2007). The bilateral superior temporal region plays a core role in the perception of both affective and linguistic prosody, as reported by most reviewed studies, and in integrating auditory and visual cues (Robins et al., 2009).

**Table 6.**
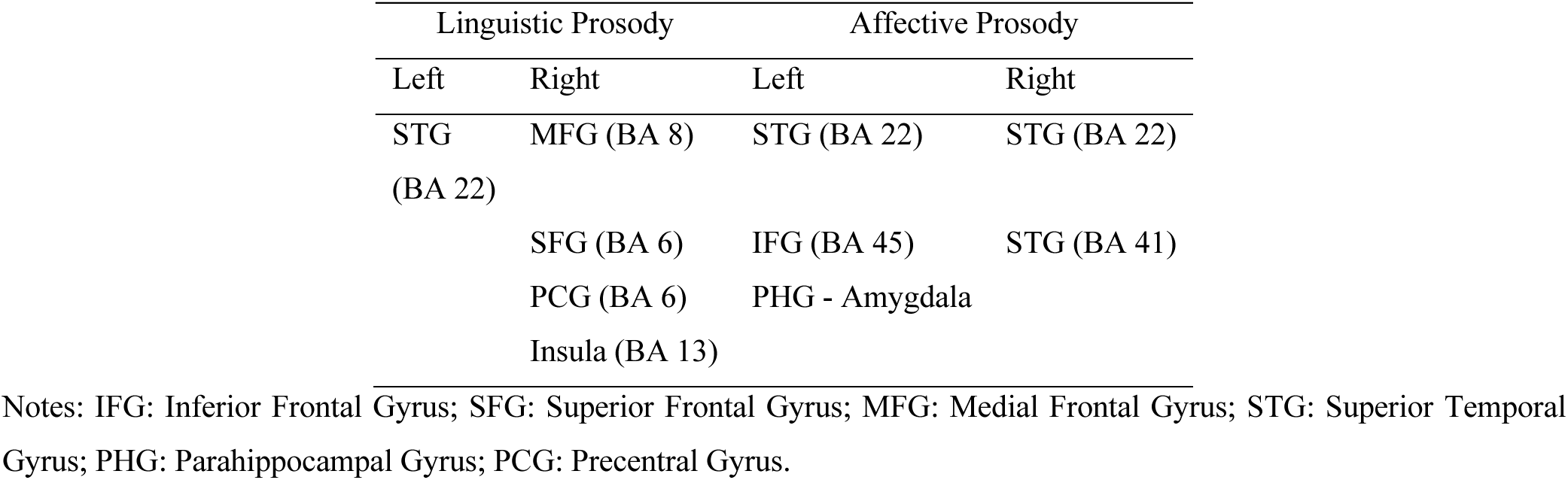
Peak activations shown after cluster analysis.

Moreover, there has been substantial evidence on the effects of subcortical structures that uniquely contribute to expressing emotions through prosody, such as the amygdala (Frühholz & Grandjean, 2013), which do not typically participate in the cortical organization of phonemes. The amygdala, part of the limbic system, connects the emotional and motivational processes, working in tandem with other brain regions to shape our memories, responses, feelings, and speech (Schirmer et al., 2008). As a critical hub for emotion detection, the amygdala shows asymmetrical activation in imaging studies when anger, fear, and, generally, emotions are triggered (Brierley et al., 2002; Schneider et al., 1997). Similarly, the insula and the striatum are also activated in the perception of emotions in patients with social phobias (Quadflieg et al., 2008). Consequently, the distinction between linguistic and affective prosody is controversial as the relationship of these two types of prosody to specific brain regions remains debated and in terms of function as prosody typically conveys both linguistic and affective meanings.

To account for these controversies, we had proposed two competing hypotheses: the *biological hypothesis* and the *linguistic hypothesis*. The *biological hypothesis* proposes that affective prosody is predominantly right-lateralized, whereas linguistic prosody should be predominantly left-lateralized. Our ALE meta-analysis showed that although affective prosody has primary activations in the right hemisphere left-lateralized areas are also active. Specifically, suitable hemisphere regions, particularly the left and right superior temporal gyrus (STG) and left inferior frontal gyrus (IFG), were activated during tasks involving affective prosody perception and production. The biological hypothesis is challenged by these findings, showing that while the right hemisphere remains engaged, left-hemisphere frontotemporal regions, common in language, are also active, suggesting a bilateral neural activation. These findings suggest that affective prosody involves in its production.

*The linguistic hypothesis* suggests that human prosodic expression is primarily integrated into the language system. Mainly, the affective prosody, to the extent that it is interlinked with the linguistic message, contains grammaticalized elements and, therefore, is regulated by language-related brain areas. This integration suggests that the prosodic aspects of language would follow standard linguistic pathways as predicted by, for example, the dual stream model of speech perception (Hickok & Poeppel, 2007). This hypothesis is not confirmed because there are active right hemisphere areas and peak activations in subcortical structures, such as the amygdala. These suggest a complex link between the affective system and the dual stream speech perception model (Hickok & Poeppel, 2007). Linguistic prosody showed strong activations in the BA22 of the left superior temporal gyrus (STG), associated with Wernicke’s area. The role of BA22 is critical for individual word understanding, sound intensity, and pitch discrimination. Sound intensity and F0 are critical for producing and perceiving lexical stress (Arvaniti, 1992; Beckman, 1986; Botinis, 1989). There are also activations in the SFG (BA 6), which is part of the pre-motor cortex and is involved in motor sequencing (Schubotz & von Cramon, 2001), movement planning (Jerjian et al., 2020), and speech articulatory programming (Flinker et al., 2015). BA8 is another area in the prefrontal cortex, responsible for the voluntary control of eye movements (saccades). This is critical as there seems to be a direct connection between pupil dilation and cognitive effort during listening and sentence planning tasks (Sevilla et al., 2014) (for a discussion, see Johnsrude and Rodd (2016)). Also, this finding corroborates an assumption about a connection between the eye and eyebrow movement, and prosody has been well reported in linguistic studies of speech prosody (Paulmann et al., 2012) and even in inner speech (Ashby & Clifton Jr, 2005).

Finally, the ALE meta-analysis shows that prosody involves more robust activation in brain areas related to social communication and interaction activations. Specifically, BA13 linked to linguistic prosody is interlinked to the social aspect of speech production being, which can explain why BA13, an area of the right insula, shows strong peak activations to linguistic prosody (Zinchenko & Arsalidou, 2018). These findings suggest that prosodic functions can correspond to an interconnected network of brain areas associated with linguistic and non-linguistic functions, unlike the production of phonemes. For example, other activations with cortical and subcortical hubs (e.g., for emotions and social cognition) determine the distinction of prosody functions. Thus, neither the linguistic nor the biological hypothesis can account for the localization problem of prosody. Affective and linguistic prosody are not isolated from each other but instead are speech expressions that convey together linguistic and emotional meanings that are interlinked both in the expression and in the underlying cognitive control, such as the connections between language and brain areas involved in social cognition and subcortical structures associated with the amygdala.

Overall, the evidence from the ALE meta-analysis shows substantial overlap in the linguistic and affective prosody concerning the brain areas involved. Likewise, these two types of prosody are expressed in speech production by shared acoustic features, such as the fundamental frequency, duration, and intensity. In other words, there are strong similarities between affective and linguistic prosody in terms of brain activations and expressive mechanisms; thus, a reasonable proposal is that they share the same prosodic mechanism. This led us to view them within a unified expressive framework, *the prosodic blending hypothesis*, which suggests that affective and linguistic, including social and other meanings are blended through selectively combined and coordinated neural activations and manifested through coordinate speech and articulatory commands. Unlike the linguistic hypothesis, however, blending provides the possibility for independent biological effects during emotional production that are integrated into the expressive mechanisms dominated by language. So, like the linguistic hypothesis, the blending hypothesis argues that there is no pure separation of emotions from of other types of information when language is involved.

The blending hypothesis addresses the interface of these two types of information. Language expressions interface primarily with linguistic information, processed mainly by the language areas in the left hemisphere; non-linguistic information, such as emotions, gaze, hand gestures, body language, and sensory data, is processed in other areas, including the right hemisphere, sensory cortices, and limbic system. The different information streams are integrated into the prefrontal cortex, temporal-parietal junction, and anterior temporal lobe, allowing the brain to create a unified understanding of meaning. Therefore, prosody interactions activate between the different brain regions, enabling the knowledge of spoken words and the general meanings expressed from the context, such as emotional activations in the amygdala. Also, the hypothesis accounts for the bilateral activations that were observed in this ALE meta-analysis, suggesting that such bilateral activations do not simply appear early due to speech perception. Instead, they are core components throughout the prosodic expression. During communication, prosodic domains express social, linguistic, and affective meanings.^7^

Thus, prosody does not differ from the other aspects of speech and grammar. For example, there is no reason to split speech sounds into affective and linguistic speech sounds, but rather argue that the segmental structure expresses both linguistic, social, and affective meanings, indicating aspects of speakers’ emotions, social and geographic identity, style, and overall communicative mechanisms. This research area has been explored in detail in our previous research (Themistocleous, 2016a, 2017a, 2017b; Themistocleous et al., 2022). Similarly, the lexicon is not viewed as social, linguistic, and affective. It is part of the language and expresses different meanings.

Therefore, the blending hypothesis addresses the interface of these two types of information. The hypothesis predicts prosody production consolidates meanings from dissimilar sources, such as linguistic, social, emotional, and conversational. It postulates that prosody should not be viewed in terms of a binary distinction between affective and linguistic prosody but rather as a unified system of gradient expressions, which is based on social and linguistic conventions but draws upon biological roots in not a dissimilar way as other domains of language, which too participate in emotions, social expression, interaction, and linguistic expression. Finally, the hypothesis suggests that prosody blends various meanings during communication. In fact, following Ohala (Ohala, 1994, 1996) and Gussenhoven (2004b), we can consider that prosody can express grammaticalized and conventional meaning deriving from biological or social functions.

### Limitations and future directions

This review included both production and perception studies in the analysis of linguistic and affective prosody. This approach has the advantage that it reports all prosody related activations in the brain whether this report prosody or perception. However, one limitation of this method is that most studies rely on perception-based experiments, so there is a great probability to report activations related to perception. Second, we have not studied individual prosodic phenomena, such as pitch accents and studies in general focus on the overall tunes, such as in statement and questions. There are more EEG studies reporting on those aspects of prosody, which were not included in the ALE meta-analysis study as they did not match our inclusion criteria (Friederici, 2017). Therefore, the analysis showed the wider activations related to linguistic and affective prosody rather than testing specific activations for specific prosody phenomena. For studying specific phenomena, currently the best approach is to manually compare the individual studies given the limited number of studies reporting such phenomena and the variability of tests and research questions. Also, the relationship of linguistic prosody to meaning requires further definition to explain how social, affective, and other meanings are produced and perceived. Our future will aim to explain the interfaces that lead from the neural activity to acoustic production and during perception from the signal to decoding the various aspects of the signal. Prosody plays a leading role in these processes.

### Data Availability Statement

The data supporting of this study are available in the Open Science Framework (OSF) at https://osf.io/vh7bg/?view_only=f22bd8f727e948c58dee230e74e3a997. The data include all relevant materials, analyses, and supplementary information necessary to replicate the study results.

---

**Appendix 1.**
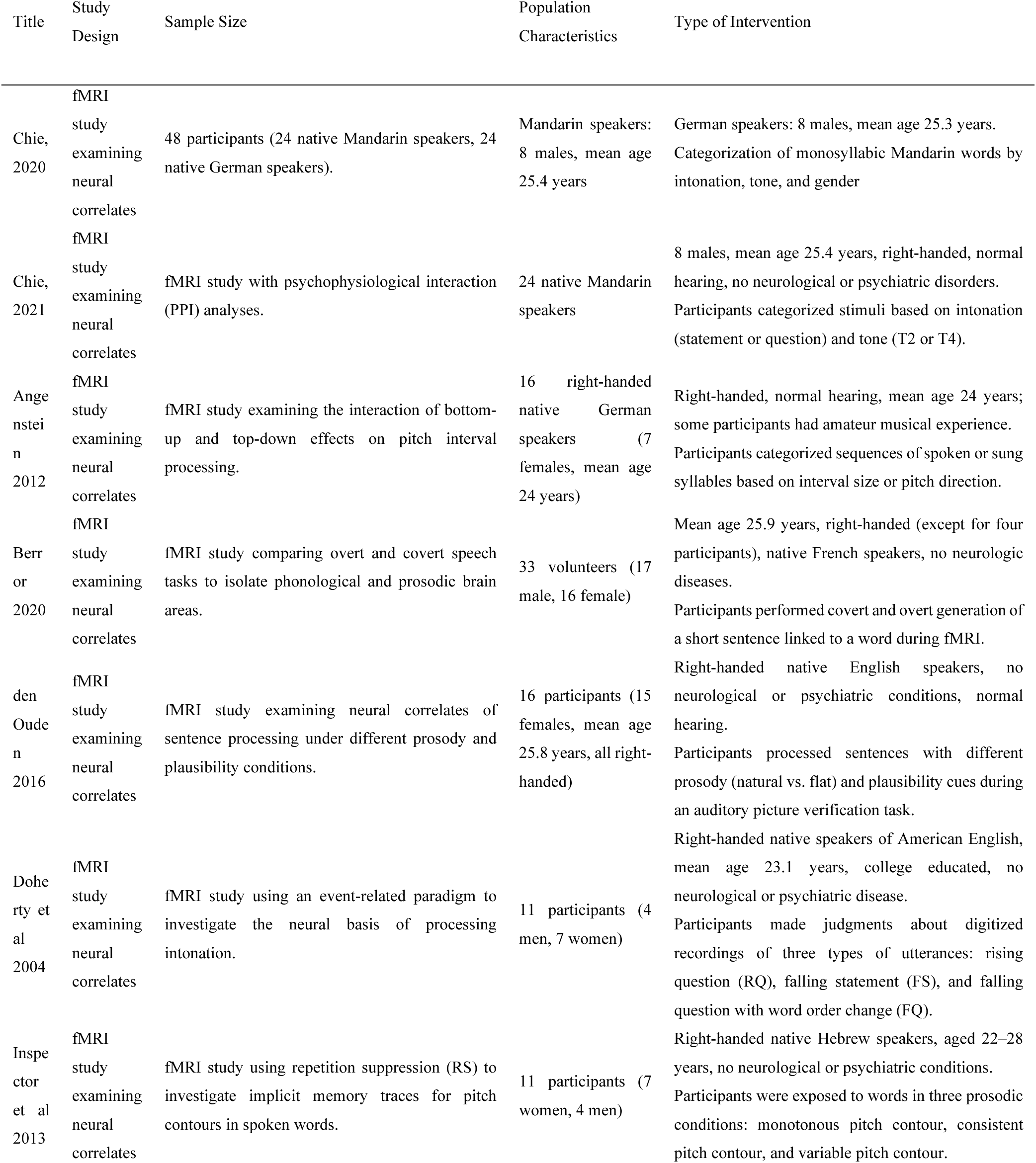
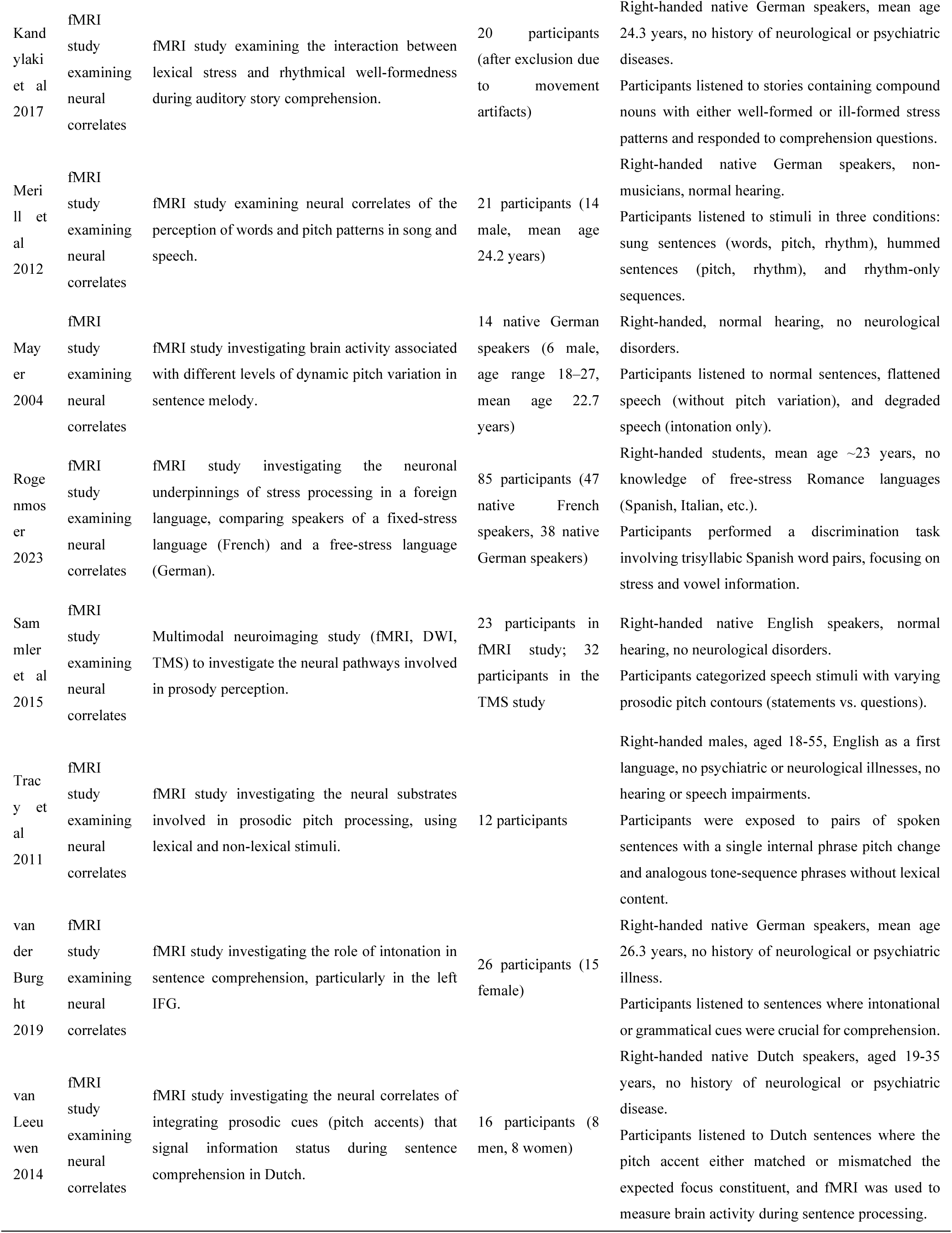
Study Design, Sample Size, Population Characteristics, and Type of Intervention of Linguistic Prosody studies.

**Appendix 2.**
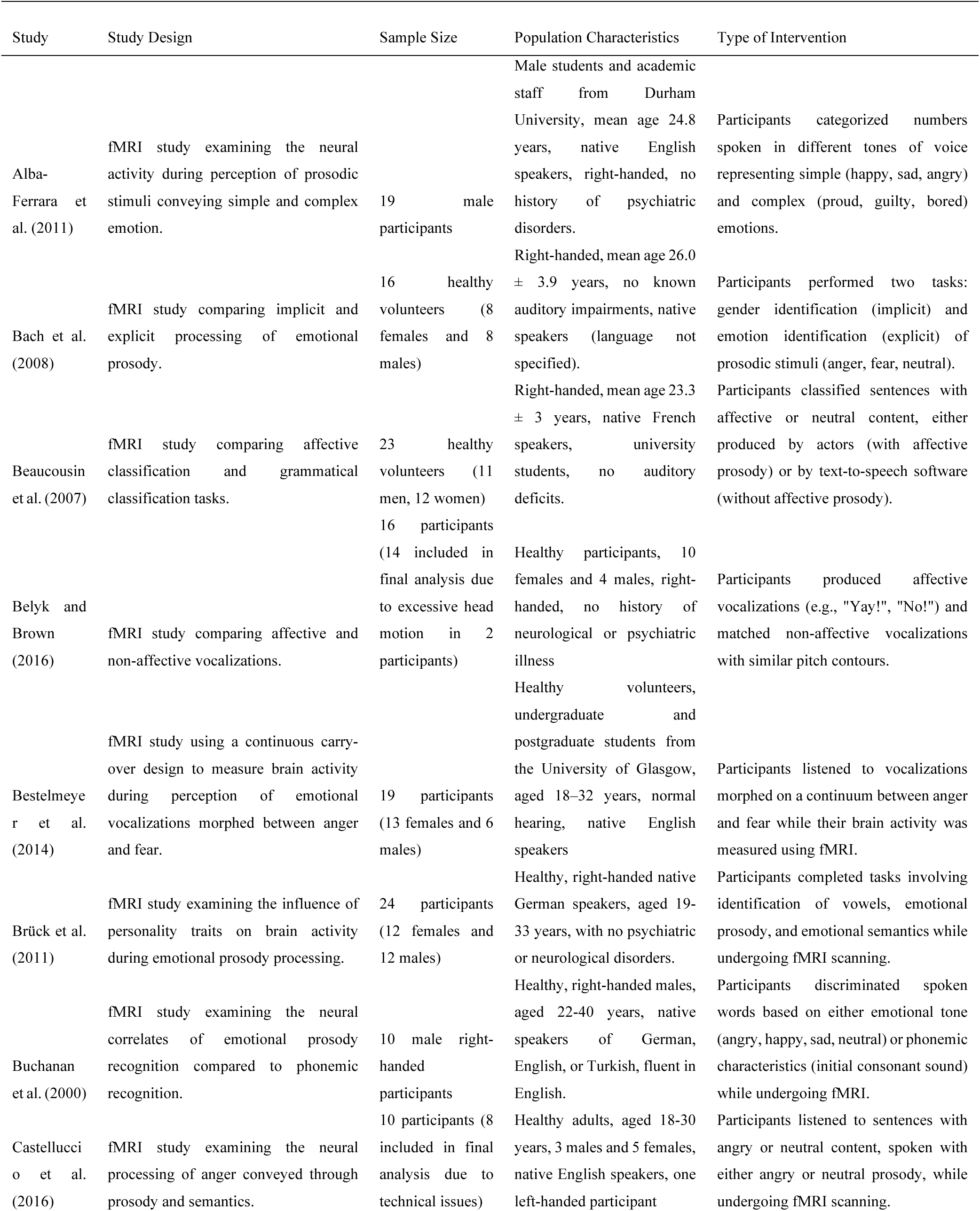
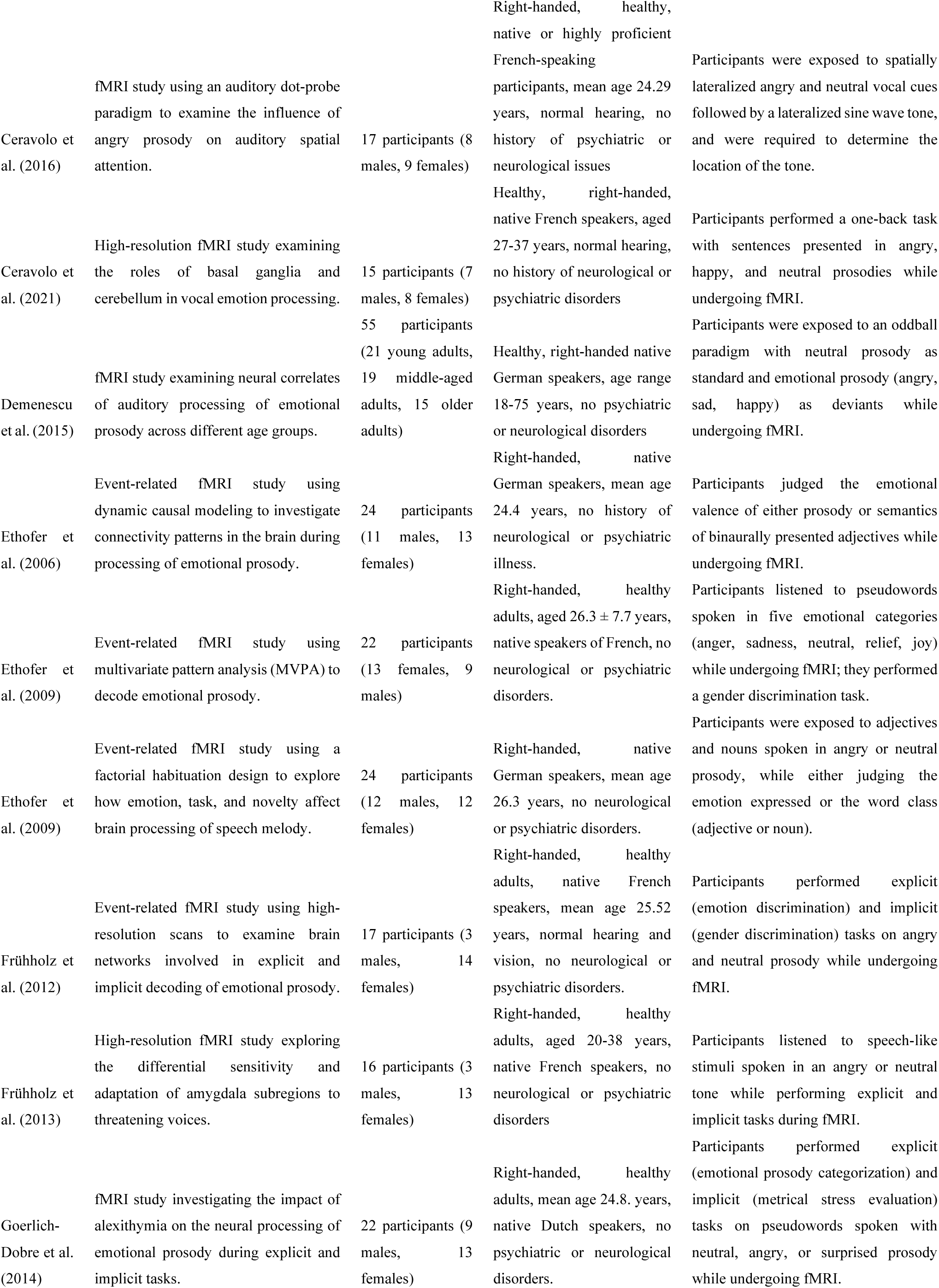
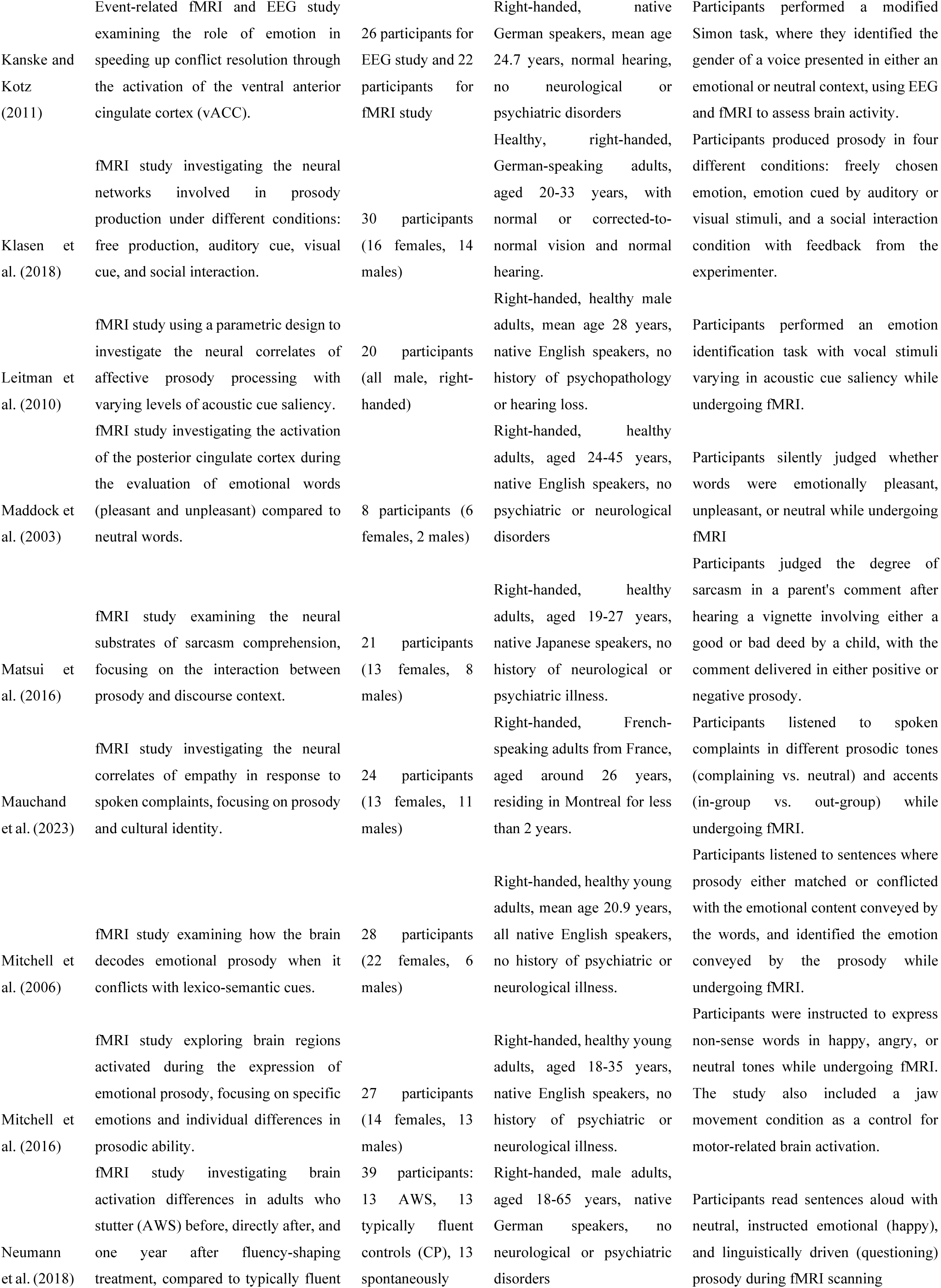
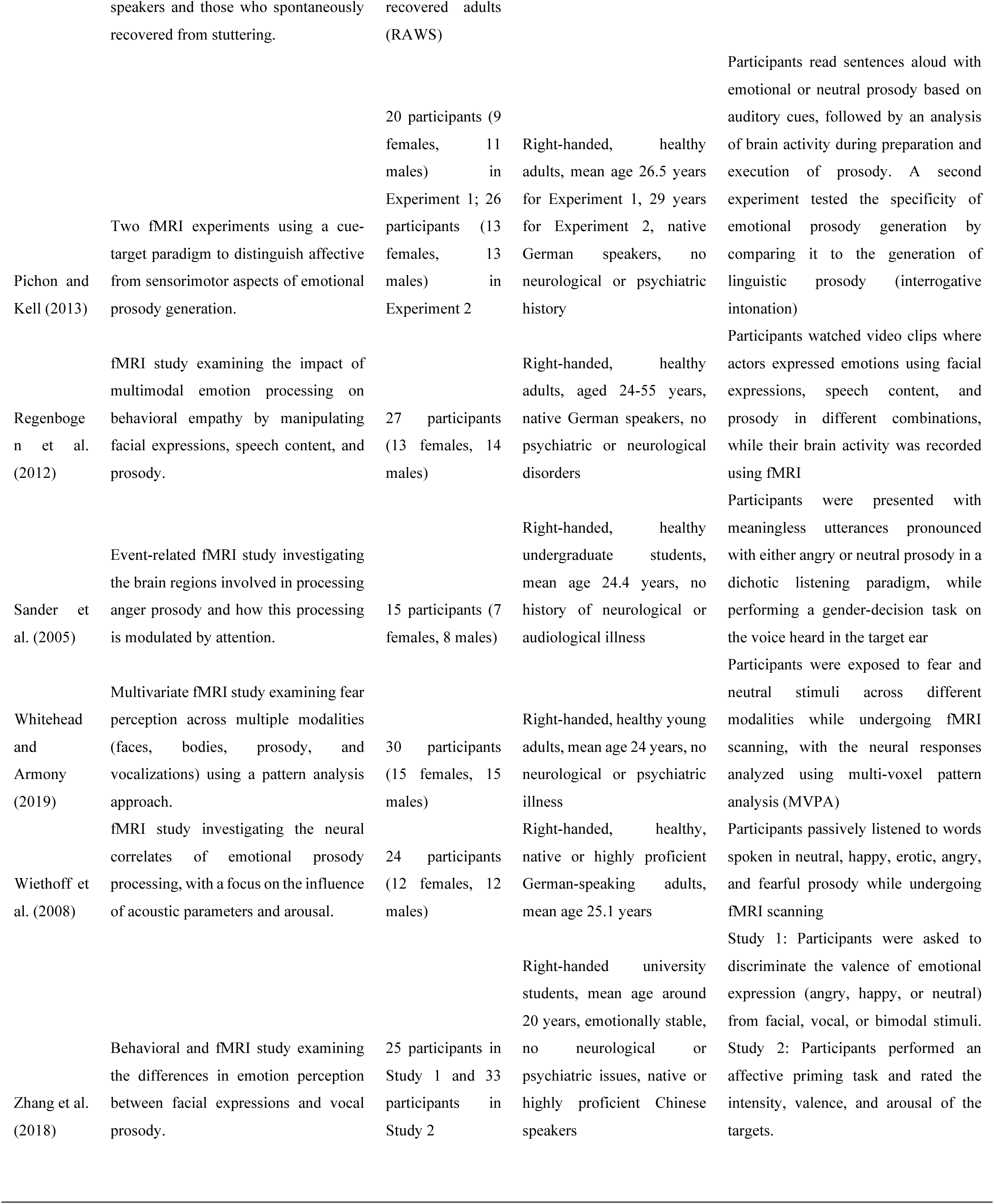
Study Design, Sample Size, Population Characteristics, and Type of Intervention of Affective Prosody studies.

## Footnotes

1 Grammaticalization typically describes the process where a content word gradually loses its lexical meaning and acquires more grammatical roles. A typical example is the Old English verb *willan* “I want to”, which lost its lexical meaning over the years and in its Modern English form “will” (as in “I will go”), indicates future tense. However, compare this to the Norwegian *vil* “will”, which still retains lexical and grammatical meanings: *Jeg vil ha en kake* “I want to have a cake.” vs. *De vil like det* “they will like it” or the “I am going to” as in I am going to the partk vs. I am going to read it Traugott, E. C., & Heine, B. (1991). *Approaches to grammaticalization*. Benjamins.

2 Similar functions were proposed by Trager, G. L., & Smith, H. L. (1951). *An outline of English structure*. Battenburg Press. who distinguished between Pitch Phonemes and Juncture Phonemes (melodic movements at contour ends) in American prosody while Bolinger, D. (1970). Relative Height. In L. Pierre, G. Gaure, & A. Rigault (Eds.), *Prosodic Features Analysis. Montreal: Didier* (pp. 107-125). Didier. (Reprinted from Bolinger, Dwight (1972), Intonation. Harmondsworth: Penguin. 137-153), Bolinger, D. (1982). Intonation and its Parts. *Language*, *58*, 505-533., Bolinger, D. (1983). Intonation and Gesture. *American Speech*, *58*(2), 156-174. who differentiated between accent (distinct tonal patterns accompanying prominent syllables) and intonation (distinct melodic movements at contour boundaries). Research from the Netherlands at the Institute of Perception Research (IPO), ’t Hart, J., Collier, R., & Cohen, A. (1990). *A Perceptual Study of Intonation. An Experimental-Phonetic Approach to Speech Melody*. Cambridge University Press. distinguished intonation into prominence-lending and non-prominence lending tonal movements, that roughly corresponds to the cumulative and delimitative functions.

3 The *F0* refers to the lowest frequency of a periodic waveform and is typically perceived as the pitch of a sound. In speech, the *F0* represents the frequency of vibration of the vocal cords during voiced segments, such as vowels. *F0* is measured in Hertz (Hz), indicating the number of vibrations (or cycles) per second of the vocal cords.

4 Despite a lot of research there is little evidence for post-nuclear pitch accents, due to tonal neutralization phenomena that manifest after a nuclear pitch accent; stresses are still being produced and perceived in post nuclear position but duration and intensity seem to play a more definite role in the actualization of those stresses Ladd, D. R. (2008). *Intonational phonology* (2 ed.). Cambridge University Press..

5 A mid (M) tone proposed by Liberman, M. (1975). *The intonational system of English* [PhD Thesis, Massachucetts Institute of Technology]. Massachucetts. is generated in Pierrehumbert, J. B. (1980). *The phonetics and phonology of English intonation* by combining H and L.

6 Although there is an effort towards standardization of the pitch accents including the role of the downstepped pitch accents !H*; there is some variation in the ToBI inventory and how it is implemented within and between languages.

7 Arguing that in an experiment would elicit only linguistic information, void of emotions and vice-versa (e.g., in non-word production) would elicit only emotional information, is highly unlikely. Emotions are likely triggered by several factors, such as stress from the experimental setting, the design, and the MRI scanner and other causes present during experimental testing. Similarly stressful and emotional factors can be triggered in real-life scenarios, such as a friendly conversation by varied factors; arguably, even a neutral emotion is an emotional state. Consequently, prosody is an expressive linguistic, social, and affective function that integrates with language, emotion, and social cognition.

